# Plasma microbiome in COVID-19 subjects: an indicator of gut barrier defects and dysbiosis

**DOI:** 10.1101/2021.04.06.438634

**Authors:** Ram Prasad, Michael John Patton, Jason L Floyd, Cristiano Pedrozo Vieira, Seth Fortmann, Mariana DuPont, Angie Harbour, Chen See Jeremy, Justin Wright, Regina Lamendella, Bruce R. Stevens, Maria B. Grant

## Abstract

The gut is a well-established route of infection and target for viral damage by SARS-CoV-2. This is supported by the clinical observation that about half of COVID-19 patients exhibit gastrointestinal (**GI**) symptoms. We asked whether the analysis of plasma could provide insight into gut barrier dysfunction in patients with COVID-19 infection. Plasma samples of COVID-19 patients (n=30) and healthy control (n=16) were collected during hospitalization. Plasma microbiome was analyzed using 16S rRNA sequencing, metatranscriptomic analysis, and gut permeability markers including FABP-2, PGN and LPS in both patient cohorts. Almost 65% (9 out 14) COVID-19 patients showed abnormal presence of gut microbes in their bloodstream. Plasma samples contained predominately *Proteobacteria, Firmicutes,* and *Actinobacteria*. The abundance of gram-negative bacteria (*Acinetobacter, Nitrospirillum, Cupriavidus, Pseudomonas, Aquabacterium, Burkholderia, Caballeronia, Parabhurkholderia, Bravibacterium,* and *Sphingomonas*) was higher than the gram-positive bacteria (*Staphylococcus* and *Lactobacillus*) in COVID-19 subjects. The levels of plasma gut permeability markers FABP2 (1282±199.6 *vs* 838.1±91.33; p=0.0757), PGN (34.64±3.178 *vs* 17.53±2.12; p<0.0001), and LPS (405.5±48.37 *vs* 249.6±17.06; p=0.0049) were higher in COVID-19 patients compared to healthy subjects. These findings support that the intestine may represent a source for bacteremia and may contribute to worsening COVID-19 outcomes. Therapies targeting the gut and prevention of gut barrier defects may represent a strategy to improve outcomes in COVID-19 patients.

## Introduction

More than 2.8 million deaths related to COVID-19 have been reported worldwide, while this number is still increasing even after more than 18 months since the diagnosis of the first COVID-19 case. According to a report published by the CDC, between the March 2020-April 2021 in the United States, the overall number of COVID-19 case are higher in female subjects (52.2%) than male (47.8%), while the mortality rate is higher among males (54.3%)^1^. The clinical manifestation of COVID-19 is more severe in patients with pre-existing and ongoing medical conditions including cardiovascular diseases, cancer, and diabetes^2–9^. Apart from respiratory symptoms, a large number of COVID-19 patients experience gastrointestinal (GI) symptoms including nausea, fever, pain, and diarrhea. Most common GI complication is severe diarrhea^10^. During hospitalization, critically ill patients experience GI complications^11^. In a USA-based study, approximately 61.3% of COVID-19 patients reported GI complications, including but not limited to loss of appetite (34.8%), diarrhea (33.7%), mesenteric arterial or venous thromboembolism, and small bowel ischemia^12 13^. These GI complications were associated with longer hospitalizations^14^. In a meta-analysis of 107 studies and 15,133 patients combined, the pooled prevalence of GI complications was 10-33.4%^15–17^. Although these studies confirm GI findings and important clinical observations, they do not interrogate the pathophysiology associated with these GI complications. We asked whether COVID-19 patients demonstrated gut barrier defects and presence of a microbiome in their plasma. Our patient population were individuals admitted to the University of Alabama at Birmingham hospital (Birmingham, AL, USA) with a confirmed diagnosis of COVID-19 and nonCOVID-19 controls.

## Methods

### Study subjects

A total of 30 COVID-19 patients (P1-P30; Table 1) and 16 healthy subjects (H1-H16; Table 2) were involved in this study. During hospitalization of the COVID-19 patients at UAB hospital, blood samples were collected under sterile conditions following Institutional Review Board guidelines. Blood samples from non-COVID-19 controls were collected following routine guidelines^18^. The COVID-19 patients were classified as mild (clinical symptoms with no sign of pneumonia), moderate (fever and respiratory symptoms), severe (any of the above criteria and following respiratory distress: ≥ 30 breaths/min; oxygen saturation: ≤ 93% at rest; arterial partial pressure of oxygen/fraction of inspired oxygen: ≤ 300 mmHg; cases with chest imaging that shows lesion progression within 24–48 h > 50%), or critical (any of the above criteria and following respiratory failure, mechanical ventilation, shock, organ failure, and requires ICU care)^19^.

**Table 1:** Basic characteristics of COVID-19 patients.

| Patient's Code | Patient's Case ID | Sex | Age (Years) | BMI | COVID-19 infection | In Hospital (Days) | Diabetes | Thrombotic Events | Current Status |
| --- | --- | --- | --- | --- | --- | --- | --- | --- | --- |
| P1 | 747 | Male | 63 | 36.94 | Mild | 49 | Yes | Yes | - |
| P2 | 784 | Male | 49 | - | Moderate | 123 | Yes | Yes | - |
| P3 | 756 | Male | 81 | - | Moderate | 30 | - | Yes | Deceased |
| P4 | 729 | Male | 49 | 35.08 | Mild | 23 | Yes | - | - |
| P5 | 765 | Female | 70 | 26.99 | Moderate | 8 | - | - | - |
| P6 | 785 | Male | 76 | - | Moderate | 4 | - | - | - |
| P7 | 678 | Female | 71 | 33.38 | Moderate | 16 | - | Yes | Deceased |
| P8 | 708 | Female | 52 | 47.55 | Mild | 14 | Yes | - | - |
| P9 | 724 | Female | 46 | 33.28 | Moderate | 25 | - | Yes | Deceased |
| P10 | 700 | Male | 55 | 37.37 | Moderate | 11 | - | Yes | - |
| P11 | 703 | Male | 56 | 27.12 | Mild | 2 | - | - | - |
| P12 | 775 | Male | 69 | 30.41 | Moderate | 25 | - | Yes | Deceased |
| P13 | 690 | Female | 69 | 41.88 | Moderate | 4 | Yes | - | - |
| P14 | 685 | Male | 48 | 29.99 | Mild | 2 | - | - | - |
| P15 | 695 | Male | 64 | 28.5 | Mild | 2 | Yes | Yes | - |
| P16 | 739 | Female | 54 | 32.24 | Mild | 1 | - | - | - |
| P17 | 704 | Female | 64 | 19.74 | Mild | 1 | - | - | - |
| P18 | 761 | Female | 70 | - | Moderate | 22 | Yes | - | - |
| P19 | 798 | Male | 55 | 29.38 | Severe | 11 | - | - | - |
| P20 | 769 | Male | 51 | 28.75 | Moderate | 6 | - | Yes | - |
| P21 | 766 | Male | 67 | 37.66 | Moderate | 1 | - | Yes | - |
| P22 | 764 | Female | 62 | 46.52 | Severe | 28 | Yes | Yes | - |
| P23 | 698 | Male | 83 | 26.63 | Moderate | 18 | - | Yes | Deceased |
| P24 | 771 | Female | 63 | 30.62 | Moderate | 4 | Yes | - | - |
| P25 | 791 | Male | 68 | 27.45 | Moderate | 22 | Yes | - | - |
| P26 | 706 | Male | 89 | 26.11 | Mild | 25 | - | Yes | - |
| P27 | 720 | Female | 73 | - | Moderate | 5 | - | - | - |
| P28 | 722 | Male | 71 | 32.61 | Mild | 5 | - | - | - |
| P29 | 694 | Female | 59 | 21.14 | Moderate | 21 | - | Yes | - |
| P30 | 693 | Female | 52 | - | Mild | 1 | - | Yes | - |

Due to low bacterial yield, sample codes P2, and P8-P11 were not included in microbiome analysis.

**Table 2:** Basic characteristics of healthy subjects. Plasma samples from these healthy controls were used in gut permeability analysis.

| **Healthy control's code** | **Sex** | **Age** | **Comorbidity** | **Current Status** |
| --- | --- | --- | --- | --- |
| H1 | Female | 35 | None | Living |
| H2 | Female | 60 | None | Living |
| H3 | Male | 56 | None | Living |
| H4 | Male | 26 | None | Living |
| H5 | Female | 43 | None | Living |
| H6 | Female | 28 | None | Living |
| H7 | Male | 42 | None | Living |
| H8 | Female | 34 | None | Living |
| H9 | Female | 34 | None | Living |
| H10 | Female | 22 | None | Living |
| H11 | Female | 72 | None | Living |
| H12 | Male | 45 | None | Living |
| H13 | Male | 55 | None | Living |
| H14 | Female | 32 | None | Living |
| H15 | Female | 33 | None | Living |
| H16 | Male | 32 | None | Living |

### Limitations of the study

Due to the small volume (200-300μL) of plasma sample’s availability, two different subsets of COVID-19 patients (P1-P14) for circulating microbiome and (P15-P30) for gut permeability marker analysis were used in the study.

### Microbial DNA extraction and 16S rRNA sequencing

The frozen plasma samples were shipped to Wright Labs, LLC. for the 16S rRNA sequencing (V3-V4 region) and metatranscriptomic analysis. The microbial DNA was extracted from samples using the DNA/RNA Miniprep Kit (Zymo Research, Irvine, CA) according to the manufacturer’s protocol. After extraction, DNA purity and concentration were determined using Qubit 4 Fluorometer (Invitrogen, Carlsbad, CA) and dsDNA HS assay kit (ThermoFisher Scientific, Waltham, MA). PCR products were pooled, and gel purified on a 2% agarose gel using Qiagen Gel Purification Kit (Qiagen, Frederick, MD). After quality check using 2100 Bioanalyzer and DNA 1000 chip (Agilent Technologies, Santa Clara, CA), 16S rRNA sequencing was performed using an Illumina MiSeq v2 chemistry with paired-end 250 base pair reads as per the Earth Microbiome Project’s protocol^20^. One negative control was processed in parallel with the samples and sequenced as well.

### Bioinformatic analysis

Raw sequence data was successfully obtained and imported into Qiime2 for processing and analyses^21^. Initial quality in the form of Phred q scores was determined using Qiime2, while cumulative expected error for each position was determined with VSEARCH^22^. Based on these quality data, forward and reverse reads were truncated at a length of 250, with a maximum expected error of 0.5 within Qiime2’s implementation of the DADA2 pipeline^23^. Qiime2’s DADA2 pipeline was also used to merge forward and reverse reads and removed chimeras and assign the remaining sequences to amplicon sequence variants (ASVs). Representative sequences were used to determine taxonomic information. The full report and statistical analysis from Wright Labs, Huntingdon, Pennsylvania is available upon request.

### Alpha and beta diversity analysis

Alpha diversity was calculated by subsampling the ASV table at 10 different depths, ranging from 230 to 2300 sequences, for the Faith’s Phylogenetic Diversity^24^, Observed OTUs^25^, Pielou’s Evenness^26^, and Shannon’s Index^27^ metrics. 20 iterations were performed at each depth to obtain average alpha diversity values for the different metrics. A rarefaction plot was created with the results of this subsampling to confirm that diversity approached an asymptote and slope decreased as depth increased. Averages for the greatest depth were calculated and plotted to show each sample’s diversity.

Beta diversity analyses were conducted after the ASV table had first undergone cumulative sum scaling normalization^28^ to mitigate differences between samples based on sequencing depth. Distances between samples were calculated using the Weighted Unifrac metric^29^ based on the normalized table and rooted tree. The resulting distance matrix was visualized as a Principal Coordinates Analysis plot in R.

### Measurement of gut permeability marker FABP2

The level of fatty acid-binding protein-2 (FABP2)^30^, a marker of intestinal barrier damage, was determined by ELISA in the plasma samples using a colorimetric assay kit (#DFBP20, R&D systems, Minneapolis, MN) following the manufacturer’s protocol. The absorbance was measured at 450nm using a microplate reader, and the levels of FABP2 were calculated as per the standard curve and expressed as pg/mL.

### Enzyme-linked immunosorbent assay for measuring gut microbial peptide translocation into the systemic circulation

The level of peptidoglycan (PGN) in plasma samples was measured using a colorimetric assay kit for mouse peptidoglycan (#MBS263268, MyBioSource Inc., San Diego, CA) following the manufacturer’s protocol. The absorbance was measured at 450nm using a microplate reader and the levels of peptidoglycans were calculated as per the standard curve and expressed as ng/mL. The levels of lipopolysaccharides (LPS) were also measured by ELISA kit (#EKC34448, Biomatik, Wilmington, DE) following the manufacturer’s instruction manual. The levels of LPS were calculated by standard curve and expressed as pg/mL.

### Statistical analysis

Data were evaluated for presence of outliers and adherence to a normal distribution using GraphPad Prism, version 8.1 software. Statistical significance of normally and non-normally distributed data were assessed via student’s *t-*test and Mann-Whitney test, respectively, at p=0.05.

## Results

### Clinical characteristics of the COVID-19 patients and healthy individuals

We enrolled 30 patients confirmed to have COVID-19 infection **(Table 1).** At the time of admission to the hospital, all COVID-19 patients were experiencing nausea, myalgia, fever, diarrhea, and shortness of breath. The median age of all 30 patients was 63.3 years (range of 46-89 years) including 17 males and 13 females. Majority of the patients (17 patients) were considered to have moderate infection, while 11 patients experienced mild infection. Only 2 patients were reported as having severe COVID-19 on admission. Based on the severity of the symptoms and duration of the recovery period, the length of the hospital stay varied from 1-123 days. Thirty-four percent of patients (10 out of 30) were diabetic and 50% of patients (15 out of 30) in our cohort experienced thrombotic events. The body mass index (BMI) of 23 patients was greater than 25. Five out of 30 patients died during their hospital stay. The blood samples from healthy individuals were collected during routine health visits and individuals were free from any complications **(Table 2).**

### Laboratory findings and COVID-19 manifestation in patients

Admission laboratories for the COVID-19 cohort included complete blood count (CBC) with differential and a metabolic panel. These results are detailed in **Tables 3 and 4**, respectively. CBC results indicated decreased abundance of lymphocytes and red blood cells (RBCs) in 47.8% and 64% of COVID-19 subjects, respectively. All COVID-19 subjects (n=23) exhibited abundance of monocytes above the normal range. Neutrophils, the first-responders of bacterial infection, were at an abundance above the normal range in 58.3% of male and 45.5% of female COVID-19 subjects. Total white blood cells counts were outside the normal range in 38.5% and 41.7% of COVID-19 subjects, respectively. Lastly, 16.7% of COVID-19 subjects exhibited platelets outside the normal range.

**Table 3:** Analytical observation of immunological cell population in the plasma samples of COVID-19 patients.

| **Patient's Code** | **Basophils (%)** | **Eosinophils (%)** | **Lymphocytes (%)** | **Monocytes (%)** | **Neutrophils (%)** | **Platelets (10<sup>3</sup>/cmm)** | **RBC (10<sup>6</sup>/cmm)** | **WBC (10<sup>3</sup>/cmm)** |
| --- | --- | --- | --- | --- | --- | --- | --- | --- |
| P1 | 0.44 | 1.33 | 17.62 | 10.26 | 70.36 | 233.44 | 2.93 | 8.18 |
| P2 | 1.02 | 1.85 | 30.34 | 6.65 | 59.05 | 498.69 | 3.99 | 9.89 |
| P3 | - | - | - | - | - | - | - | - |
| P4 | 0.68 | 1.8 | 14.52 | 7.36 | 75.23 | 335.08 | 3.17 | 8.94 |
| P5 | 0.4 | 0.13 | 8.2 | 6.7 | 86 | 307.47 | 4.87 | 9.03 |
| P6 | - | - | - | - | - | - | - | - |
| P7 | 0.27 | 0.1 | 1.81 | 1.59 | 96.37 | 166.37 | 3.00 | 6.79 |
| P8 | 0.18 | 0.07 | 85.13 | 2.27 | 13.36 | 164.13 | 4.06 | 39.76 |
| P9 | 0.84 | 2.09 | 22.52 | 6.15 | 68.64 | 295.4 | 3.4 | 8.78 |
| P10 | 0.75 | 2.6 | 13.24 | 5.7 | 78.11 | 191.06 | 4.36 | 11.06 |
| P11 | - | - | - | - | - | - | - | - |
| P12 | 0.3 | 0.1 | 11.1 | 3.0 | 86 | 211.7 | 4.6 | 5.51 |
| P13 | 0.2 | - | 36.8 | 6.6 | 56 | 151.9 | 5.09 | 3.59 |
| P14 | 0.57 | 2.76 | 18.16 | 7.15 | 71.78 | 276.2 | 2.84 | 9.63 |
| P15 | 1.27 | 2.5 | 27 | 13.33 | 56.03 | 142.4 | 2.74 | 3.84 |
| P16 | 0.6 | 0.6 | 26.8 | 6.3 | 66 | 423.8 | 4.91 | 16.86 |
| P17 | 0.5 | 2.1 | 43.7 | 13.4 | 40 | 82.7 | 2.88 | 6.26 |
| P18 | 0.4 | 0.3 | 15.3 | 9.8 | 74 | 185.5 | 3.91 | 8.16 |
| P19 | 0.2 | 0.4 | 5.4 | 6.1 | 88 | 341.4 | 4.1 | 9.71 |
| P20 | 0.2 | 0.2 | 22.2 | 7.5 | 70 | - | 5.33 | 7.09 |
| P21 | - | - | - | - | - | 268.8 | 4.39 | 9.34 |
| P22 | 0.4 | 0.1 | 10.9 | 9.0 | 82 | 231.7 | 4.96 | 6.99 |
| P23 | - | 1.1 | 3.2 | 3.9 | 93 | 290.1 | 4.2 | 27.66 |
| P24 | - | - | - | - | - | 263.2 | 5.32 | 4.52 |
| P25 | 0.8 | 0.3 | 4 | 12.7 | 82 | 308.6 | 4.11 | 14.62 |
| P26 | - | - | - | - | - | - | - | - |
| P27 | 0.4 | 0.4 | 30.3 | 21.2 | 48 | 328.9 | 3.49 | 2.10 |
| P28 | - | - | 4 | 4 | 92 | 241.2 | 4.72 | 15.54 |
| P29 | 0.9 | 1.1 | 15.6 | 5.4 | 77 | 161.1 | 2.9 | 13.68 |
| P30 | - | - | - | - | - | - | - | - |

RBC: Red blood cells; WBC: White blood cells.

**Table 4:** Biochemical observations in the plasma samples of COVID-19 patients.

| Patient's Code | BNP (pg/mL) | CRP (mg/L) | D-Dimer (ng/mL) | Ferritin (ng/mL) | Glucose (mg/dL) | Hgb (gm/dL) | Lactic acid (mm/L) | LDH (units/l) | Hs Troponin-I (ng/L) | Procalcitonin (ng/mL) |
| --- | --- | --- | --- | --- | --- | --- | --- | --- | --- | --- |
| P1 | 515 | 56.27 | 1914.51 | 503.67 | 160.11 | 8.56 | 1.34 | 291 | 182.92 | 0.7 |
| P2 | 77 | 64.89 | 439.62 | 351 | 212.81 | 11.28 | 1.24 | 320.33 | - | - |
| P3 | - | - | - | - | - | - | - | - | - | - |
| P4 | - | 208.61 | - | 187 | 202.74 | 8.69 | 1.8 | - | 6.5 | - |
| P5 | 69 | - | - | - | 120.5 | 12.8 | - | - | 5.3 | - |
| P6 | - | - | - | - | - | - | - | - | - | - |
| P7 | - | 67.77 | 1298 | 1489 | 148.35 | 8.89 | 1.4 | 1636.9 | 28.17 | 0.08 |
| P8 | 25 | 25.12 | 308.38 | 108 | 134.3 | 10.39 | 0.8 | 346.33 | 3.33 | 0.02 |
| P9 | 53 | 138.16 | 188.69 | 4552 | 123.73 | 9.91 | 1.84 | 503 | 8.57 | 0.8 |
| P10 | 136 | 431.79 | 1744.77 | 1462 | 114 | 13.93 | 2.76 | 2221 | 98.27 | 26.46 |
| P11 | - | 115.98 | 176.82 | 292 | 195.5 | 12.9 | - | 221 | - | 0.07 |
| P12 | 22 | 26.1 | 568.57 | 2105 | 175 | 15.4 | - | 331 | 24.3 | 0.09 |
| P13 | 51 | 59.01 | 577.28 | 379 | 137.69 | 7.91 | 0.79 | 399 | 5.1 | 0.12 |
| P14 | - | 25.92 | 218.04 | 477 | 104.39 | 15.96 | - | 273 | - | 0.07 |
| P15 | - | - | - | - | 78 | 8 | - | - | - | - |
| P16 | - | - | - | - | 118 | 14.3 | - | - | - | - |
| P17 | - | - | - | - | 262 | 9.5 | - | - | - | - |
| P18 | - | - | - | - | 111 | 10.2 | - | - | - | - |
| P19 | - | - | - | - | 143 | 11.8 | - | - | 2.3 | - |
| P20 | - | - | - | - | 220 | 14.0 | - | - | - | - |
| P21 | - | - | - | - | 100 | 12.7 | - | - | - | - |
| P22 | 88 | 130.22 | 410.68 | 63 | 167 | 13.1 | 1.8 | 272 | 4.0 | 0.07 |
| P23 | 551 | 297.31 | 9842.0 | - | 207 | 12.3 | - | - | 2865.8 | - |
| P24 | - | - | - | - | 420 | 12.9 | - | - | - | - |
| P25 | - | - | - | - | 301 | 12.2 | - | - | - | - |
| P26 | - | - | - | - | 99 | - | - | - | - | - |
| P27 | - | 26.67 | 632.68 | - | 113 | 9.5 | - | 303 | - | - |
| P28 | - | 20.76 | 414.04 | - | 113 | 14.5 | - | 280 | - | - |
| P29 | - | - | - | - | 99 | 10.0 | - | - | - | - |
| P30 | - | - | - | - | - | - | - | - | - | - |

BNP: Brain natriuretic peptide; CRP:C-reactive protein; Hgb: Hemoglobin; LDH: Lactate dehydrogenase.

Biochemical evaluation of these patients was performed as seen in **Table 4**. Of COVID-19 positive subjects, 100% of those with BNP out of range (n=3) were male of which one subject exhibited BNP between 100-200 pg/mL, likely indicative of compensated congestive heart failure (CHF), and two subjects with BNP >400 pg/mL, indicating likely moderate to severe CHF. All subjects exhibited CRP levels greater than normal of which four male subjects (44%) and two female subjects (33%) exhibited CRP >100 mg/L which is associated with severe inflammation such as sepsis. Six male (75%) and six female (100%) subjects exhibited D-dimer levels higher than the normal range, indicating activation of the procoagulant and fibrinolytic systems. Five male (71.4%) and three female (60%) exhibited ferritin levels greater than the normal range of which two male (40%) and two female subjects (67%) had ferritin levels >1000 ng/mL which can be associated acute or chronic inflammation. Twelve male (80%) and 11 female (91.7%) exhibited fasting glucose levels greater than the normal range. Nine male (64.3%) and 8 female (66.7%) subjects exhibited hemoglobin levels below the normal range. Six male (85.7%) and six female (100%) subjects exhibited LDH levels greater than normal. Of male subjects, four subjects (80%) exhibited troponin-I levels above the normal range and one subject (20%) had levels below the normal range. Of female subjects, one (16.7%) exhibited troponin-I levels greater than the normal range. Troponin-I levels greater than the normal range suggest myocardial injury. Three male (60%) and three female (60%) subjects exhibited procalcitonin levels greater than the normal range with one male (33.3%) and one female (33.3%) subject exhibiting levels between 0.15-2.0 ng/mL and one male (33.3%) subject with a procalcitonin level greater than 2 ng/mL. Procalcitonin levels <0.15 ng/mL indicate an unlikelihood of significant bacterial infection; whereas, levels between 0.15-2.0 ng/mL do not exclude the possibility of an infection, and levels >2.0 ng/mL are highly suggestive of a significant bacterial infection. Reference values for laboratory measurements are provided in **table 5.**

**Table 5.**
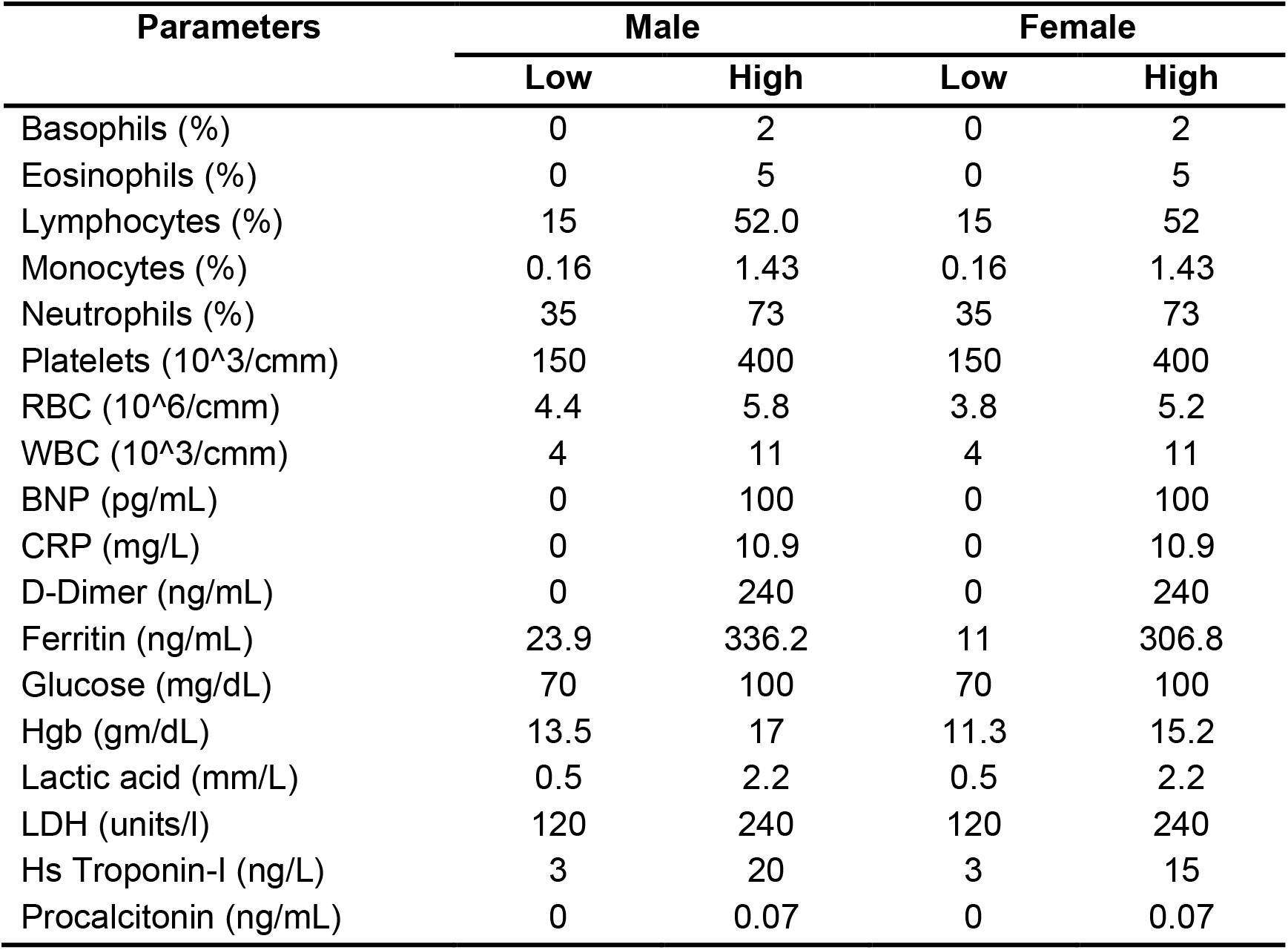
Reference values for CBC with differential and biochemical panel in healthy individuals.

### Presence of gut microbial abundance in the blood of COVID-19 patients

Plasma samples were obtained under sterile conditions and evaluated for the presence of bacteria. Specifically, the taxonomic units, distribution of abundances, and alpha diversity were measured. Alpha diversity, a representation of the total microbial population in the sample, was assessed using Pielou’s Evenness, Faith’s Phylogenic Diversity, Observed Features Metrics, and Shannon’s Index, **(Fig.1A-D).** A total of 152,536 sequencing reads were obtained from 14 COVID-19 plasma samples. 16S rRNA sequencing data suggests that 65% (9 out of 14 patient samples) yielded a strong bacterial signal. Alpha diversity revealed that the plasma microbiome for each patient exhibited unique evenness and richness. However, notable differences were observed in the Pielou’s, Faith’s, Observed, and Shannon index between samples. Beta diversity was determined using principal coordinate analysis (PCoA) **(Fig.1E)**. Overall, the plasma microbiome community was not different between the COVID-19 samples.

**Figure 1:**
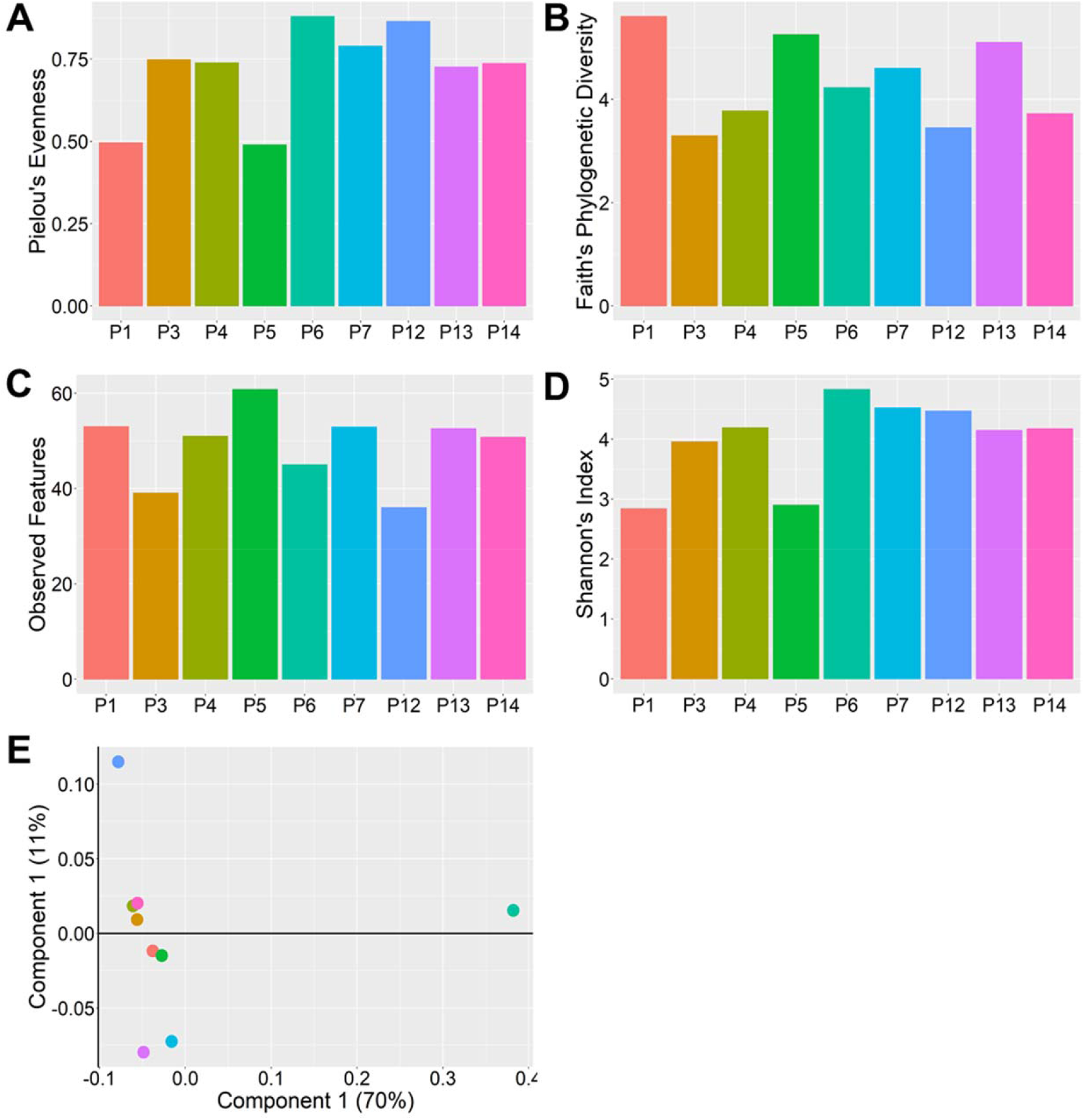
16S rRNA analyses in the plasma of COVID-19 patients. Alpha diversity was measured by observed species richness within the samples **(A-D)**. 2D principal coordinates analysis (PCoA) plots of weighted UniFrac distance reveal no difference among the patients**(E)**.

### Phylogenic differences in plasma microbiome in the COVID-19 plasma samples

The dysbiosis index is a PCR-based assay and was performed to quantify the abundance of bacterial groups in the given samples. As shown in **Fig.2A**, a dysbiosis index was determined in the plasma of all the COVID-19 samples. The relative abundance of microbial composition in the COVID-19 samples is shown in **Fig.2B**. Three major phyla (*Proteobacteria, Firmicutes*, and *Actinobacteria*) were identified in all 9 samples. Patient 6, however, exhibited abundance of unidentified bacteria greater than all other subjects. At the phylum level, the enrichment of *Proteobacteria* was highest in all samples ranging from 22%-91%, followed by *Firmicutes* (10%-71%), and *Actinobacteria* (6%-27%). *Bacteroidetes* was present in a very low percentage. *Firmicutes* abundance in P7 and P12, two of those which died during hospitalization, was low, suggesting plasma abundance of *Firmicutes* may be a prognostic marker of COVID-19 severity.

**Figure 2.**
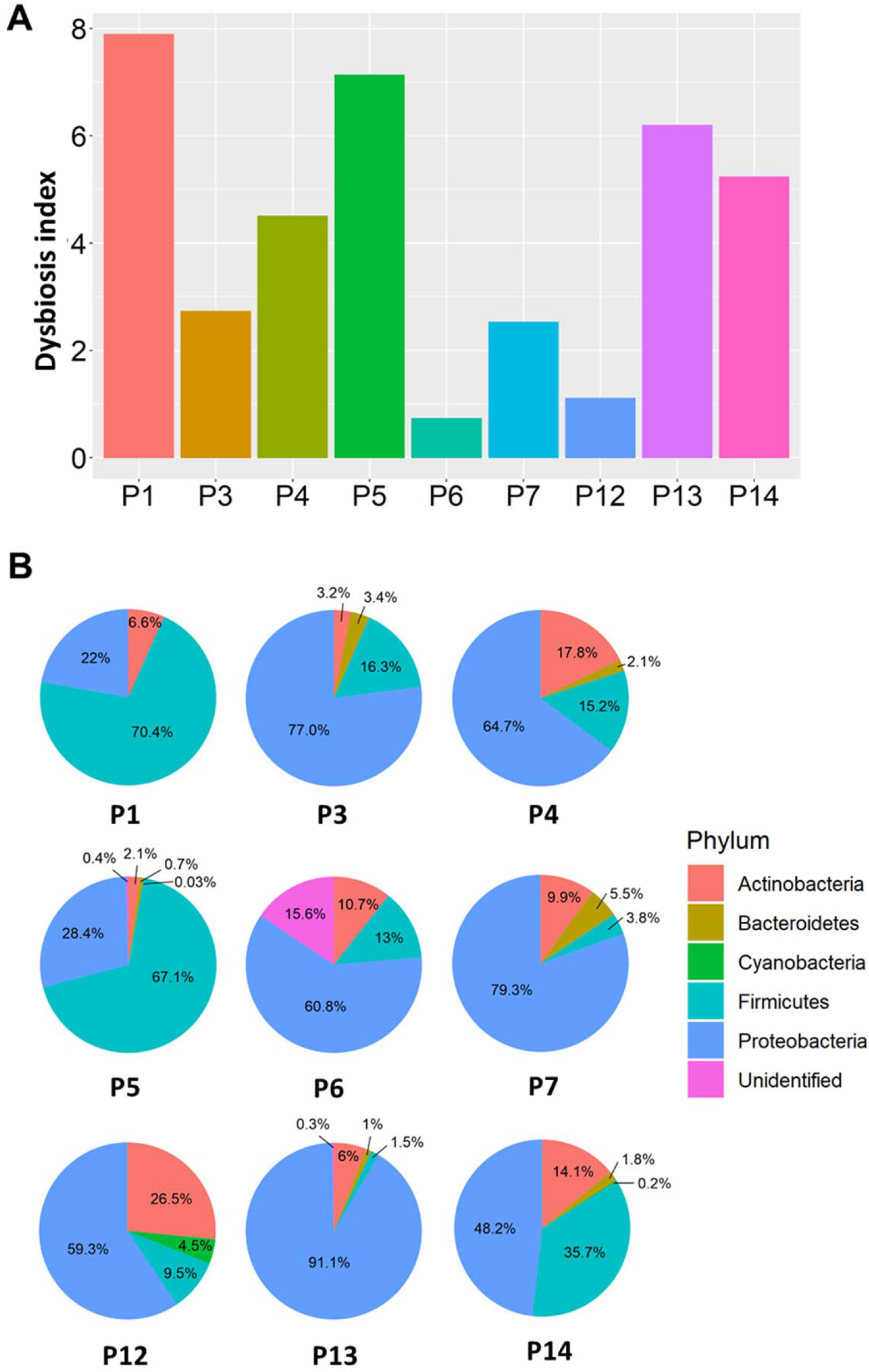
Pathogen dysbiosis index **(A)** and dominant phyla **(B)** were observed in COVID-19 plasma. Pie charts representing the main phyla that constitute the blood microbiome in COVID-19 samples.

### Taxonomic abundance in the COVID-19 plasma samples

Next, the abundance of each microbial population was assessed and revealed that, at the genus level (**Fig.3),** the prevalence of gram-negative bacteria (*Acinetobacter, Nitrospirillum, Cupriavidus, Pseudomonas, Aquabacterium, Burkholderia, Caballeronia, Parabhurkholderia, Bravibacterium,* and *Sphingomonas*) was higher than gram-positive bacteria (*Staphylococcus* and *Lactobacillus*) in COVID-19 plasma samples. Notably, LPS, a major cell wall component of gram-negative bacteria which contributes to the activation of inflammatory signaling pathways, was significantly increased in COVID-19 subject plasma (p=0.0049), supporting the observed increase of gram-negative bacteria.

**Figure 3:**
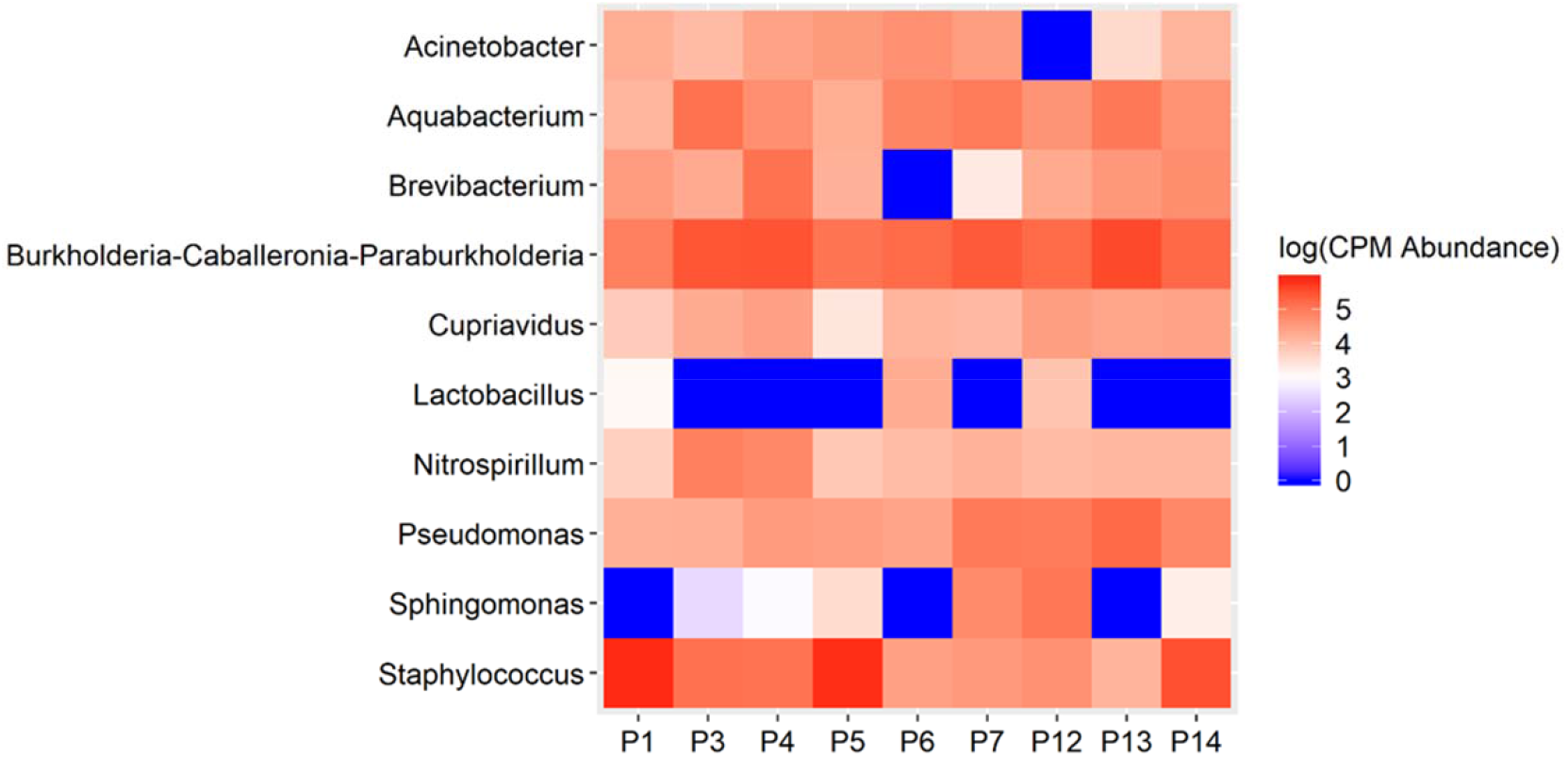
A heatmap analysis of CPM normalized counts of Metaphlan displayed differential abundances of several prominent taxa in the COVID-19 plasma samples.

### SARS-CoV-2 infections promote gut barrier defects in COVID-19 patients

The plasma microbiome arises largely as a consequence of bacterial translocation from the gut into the systemic circulation^31–36^. Compromised intestinal barriers are an important pathogenic factor and contribute to promotion of inflammation. We measured gut permeability markers in the plasma of COVID-19 and control subjects. FABP2 is an intracellular protein which is expressed specifically in intestinal epithelial cells^37^ and binds free fatty acids, cholesterol, and retinoids, and is involved in intracellular lipid transport. During mucosal damage, mature epithelial cells release this protein into the circulation^38^ and higher levels of FABP2 in the plasma are associated with gut barrier defects^30 37 39 40^. To determine the integrity of the gut barrier in COVID-19 patients, the levels of FABP2 were measured. As seen in **Fig.4A**, the levels of FABP2 were higher in the plasma of COVID-19 patients (1282±199.6 *vs* 838.1±91.33; p=0.0757) compared with healthy individuals.

**Figure 4.**
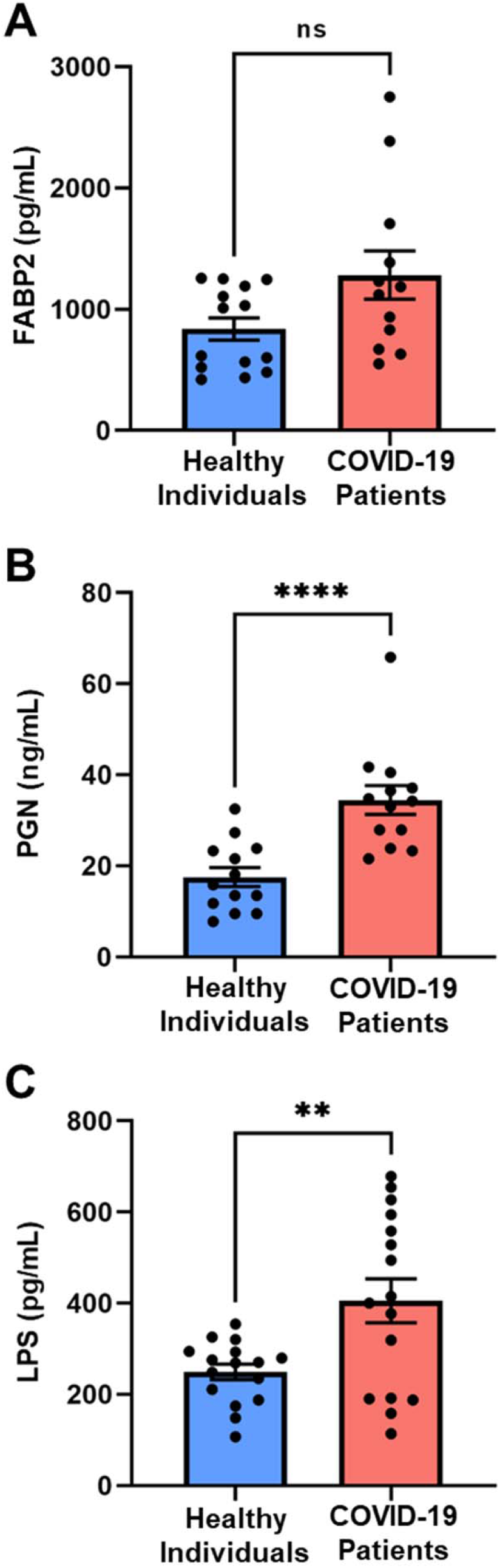
The SARS-CoV-2 infection causes gut barrier dysfunctions measured by higher levels of FABP2 **(A),** PGN **(B),** and LPS **(C)** in the plasma of COVID-19 patients. Data are presented as mean ± S.E.M. Each dot represents a sample in the cohorts. *p<0.07, **p<0.0001, ***p<0.0049.

### The increased levels of gut microbial peptides (GMPs) into systemic circulation

GMPs initiate deleterious signaling pathways and contribute to systemic inflammation^31 41–50^. To determine if gut barrier dysfunction led to translocation of GMPs into the system circulation of COVID-19 patients, we measured PGN and LPS in their plasma. Levels of PGN were nearly 2.5 times higher (34.64±3.18 *vs* 17.53±2.12; p<0.0001) in COVID-19 patients compared with controls **(Fig.4B)**. LPS, the major component of gram-negative bacterial cell walls, was found in higher levels (405.5±48.37 *vs* 249.6±17.06; p=0.0049) in COVID-19 samples compared with non-COVID-19 patients **(Fig.4C).**

## Discussion

Due to their role in regulating immune function and metabolism, gut microbes are key contributors in the maintenance of host health^51–54^. The fecal microbiota and its translocation from the gastrointestinal tract into systemic circulation has been considered as a key driver of immune response and systemic inflammation ^55–58^. Abnormal presence of gut microbes in the plasma can initiate and intensify inflammatory cascades^59^. Although systemic and local tissue inflammation is paramount in the pathogenesis of COVID-19 infection, the clinical relevance of gut microbes in the plasma remains unclear. Therefore, in this study we sought to test the hypothesis that bacterial translocation from the intestine into the systemic circulation occurs and is associated with worsened outcomes in SARS-CoV-2 infection. Increased intestinal permeability due to mucosal barrier dysfunction could result in microbial translocation. Our results support that the COVID-19 patients exhibit gut barrier dysfunction as evidenced higher levels of FABP2, PGN, and LPS **(Fig.4)** and the presence of microbes in their plasma **(Fig.1–3)**.

The duration of fecal viral shedding ranged from 1 to 33 days after symptomatic recovery of lung pathology ^60 61^. In children infected with SARS-CoV-2, rectal swabs were found positive for SARS-CoV-2 even after the nasopharynx was negative, suggesting that viral shedding from the digestive tract might be longer duration than that from the respiratory tract^62^.

During hospitalization, the fecal microbiome can be altered, thus, we selected to test the initial plasma samples of COVID-19 patients. In a small group of 9 patients, depletion of the commensal bacterium *Lactobacillus* was documented in 65% patients during COVID-19 infection. Commensal bacteria act on the host’s immune system to induce a protective response and also inhibit the growth of respiratory pathogens^63^. Heeney *et. al*. reported reduced abundance of Lactobacillus in diabetes, obesity, and cancer ^64^. Our data in **Table 1** suggests that majority of COVID-19 patients were diabetic and obese as depicted from higher BMI. The microbial abundance of *Acinetobacter, Nitrospirillum, Cupriavidus, Pseudomonas, Aquabacterium, Burkholderia, Caballeronia, Parabhurkholderia, Bravibacterium,* and *Sphingomonas*) were higher in COVID-19 patients. Sepsis is defined as a life-threatening condition in which body’s immune system damages its own tissues in response to infections ^65^. The increased abundance of Gram-positive bacteria *Staphylococcus* in the bloodstream can cause sepsis and infective endocarditis ^66^. Alhazzani et reported that most of the COVID-19 related deaths are caused by sepsis ^67^. Even after viral clearing, there was a loss of salutary species in the majority of COVID-19 patients, suggesting that exposure to SARS-CoV-2 might be associated with more long-lasting deleterious effects on the gut microbiome.

While most studies to date examine the blood metabolome, rather than the blood microbiome, we first sought to establish whether the plasma microbiome existed in COVID-19 subjects and then determine if the microbial diversity supported that the origin of these microbes was the intestine^68 69^. Results from numerous studies have linked the plasma metabolome to the gut microbiome and their implication for specific diseases^70^. Wikoff *et. al.* demonstrated that the gut microbiome dramatically influenced the composition of blood metabolites using MS-based methods and plasma extracts from germ-free mice compared with samples from conventional animals^71^. Bacterial-mediated production of bioactive indole-containing metabolites derived from tryptophan such as indoxyl sulfate and the antioxidant indole-3-propionic acid (IPA) have been identified in the plasma.

The fecal microbiome has been compared to the plasma metabolome in disease states such as ulcerative colitis where products of sphingolipid metabolism, specifically sphingosine 1-phosphate in the blood correlate with *Roseburia*, *Klebsiella*, and *Escherichia-Shigella* ^72^. Kurilshikov *et. al.* showed that the gut microbiome explained 11% to 16% of the variation in 231 major plasma metabolites^73^, highlighting its powerful impact on the host and the multidimensional interplay between gut bacteria and their ability to predict human disease or health.

Studies on the plasma microbiome are limited; however, Whittle et. al. performed a comprehensive evaluation of the blood microbiome in healthy and asthmatic individuals and found, at the phylum level, the blood microbiome was predominately composed of *Proteobacteria, Actinobacteria, Firmicutes*, and *Bacteroidetes*^74^. These key phyla detected were consistent irrespective of molecular method used for their identification (DNA vs. RNA), and were consistent with the results of other published studies^75–78^.

Studies by Serena *et. al.* demonstrate that celiac disease patients exhibit alterations in blood microbiome composition and taxonomic diversity compared to healthy subjects and they suggested that changes in the blood microbiome may contribute to the pathogenesis of celiac disease^79^. Buford et al compared microbiota profiles of serum from healthy young (20–35 years) and older adults (60–75 years). They demonstrated that the richness and composition of the serum microbiome differ between these age groups and are linked to indices of age-related inflammation such as IL-6 and TNFα^80^.

Our studies provide evidence for the loss of gut barrier function in COVID-19 subjects, however, the mechanisms responsible have not been elucidated. A role for loss of intestinal angiotensin converting enzyme 2 and a dysregulated renin-angiotensin system is plausible. SARS-CoV-2, upon entry into the host, binds to the extracellular domain of ACE2 in the nose, lung, and gut epithelial cells through its spike glycoprotein subunit S1. In a healthy gut, ACE2 serves to chaperone amino acid transporters to the gut epithelial surface. At the gut epithelial surface, ACE2 dimerizes with B^0^AT1 and then a dimer of ACE2: B^0^AT1 heterodimers activates mucosal enteroendocrine L cells to release incretins, such as GLP-1 and GIP. Incretins enter the circulation to modulate glucose homeostasis. This key regulatory pathway can be disturbed in patients with COVID-19, even nondiabetics, and might explain, in part, new onset hyperglycemia that is seen in COVID-19 patients as well as deterioration of glucose control in diabetic subjects with COVID-19 infections. Amino acid absorption in the gut, regulated by the ACE2:B^0^AT1 modulates not only tryptophan absorption but also glutamine, and tryptophan serves to activate mTOR to release antimicrobial peptides that signal to down-regulate lymphoid proinflammatory cytokines and promote tight junction formation ^81 82^. These natural defense mechanisms are disturbed by COVID-19 infection and can lead to a leaky gut.

This study has limitations including the absence of plasma microbiome samples in control subjects and the rather small size for COVID-19 subjects. Despite the limitations, we show conclusively that gut barrier leakage occurs in COVID-19 subjects. Taken together, we summarized our observations and show the presence of pathogenic bacteria in the plasma of COVID-19 subjects **(Fig. 5)**. SARS-CoV-2 infection disrupts the gut barrier and leads to elevation of systemic bacterial lipopolysaccharide and peptidoglycan and serves to enhance systemic inflammation. Therefore, leaky gut and microbial dysbiosis could contribute to cytokine storm in patients severely ill with COVID-19.

**Figure 5.**
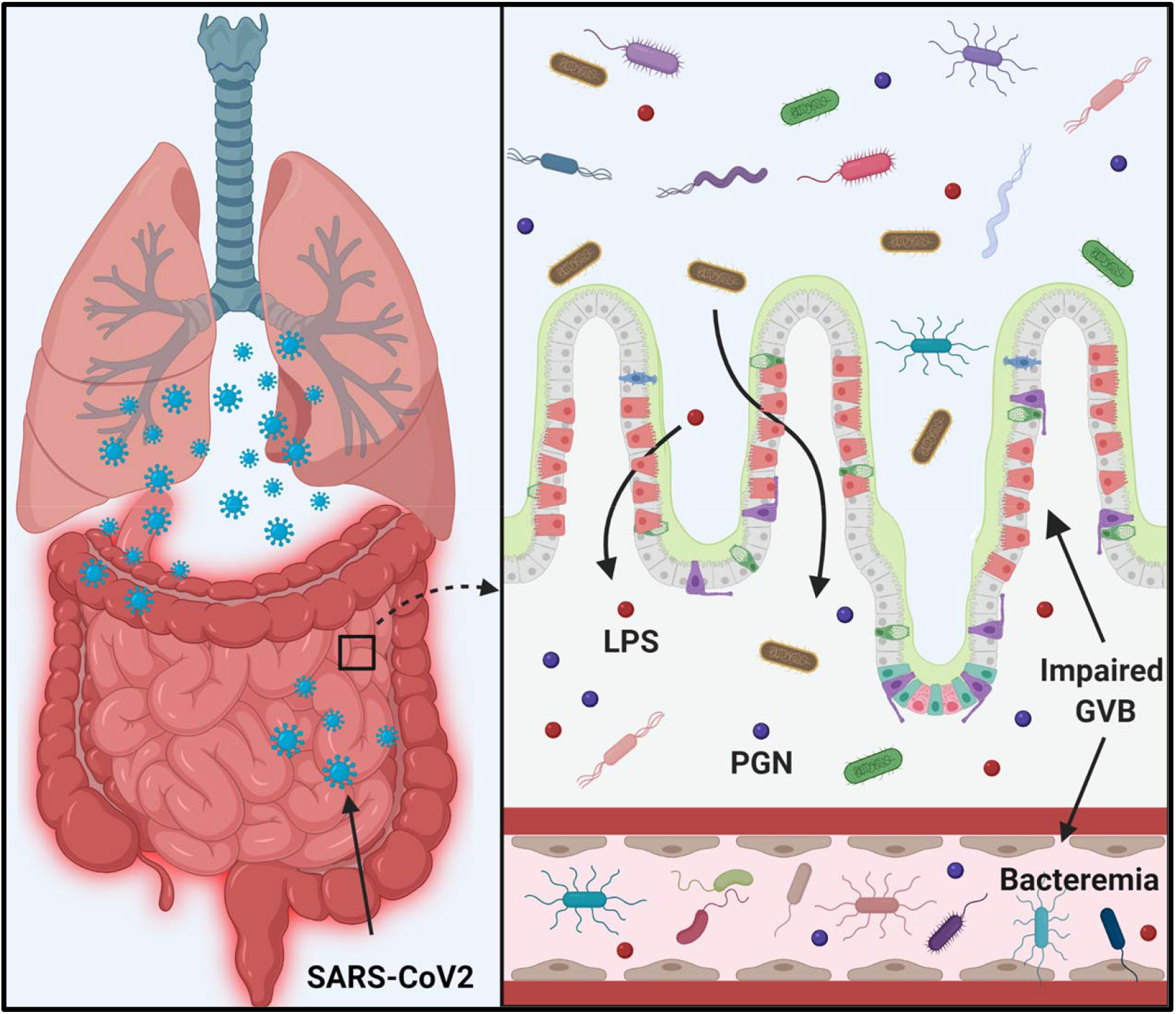
Schematic diagram representing hypothesis of COVID-19 infection promotes gut barrier defects and translocation of gut microbiome into the systemic circulation resulting worse outcomes.

## Authors Contributions

RP, MJP, and MBG conceived of the study and participated in the design. RP, MJP, SF, CPV, and MD were involved in sample collection. CSJ, JW, RL performed microbiome analysis. RP performed ELISA for gut permeability markers. RP and JLF performed data analyses and all figures, including Figure 5 with BioRender. RP, JLF, AH, and MBG wrote the manuscript. BRS provides expert guidance and critiques. All authors read and approved the final manuscript.

